# Soluble Syntaxin 3 Functions as a Transcription Regulator

**DOI:** 10.1101/162958

**Authors:** Adrian J. Giovannone, Christine Winterstein, Pallavi Bhattaram, Elena Reales, Seng Hui Low, Julie E. Baggs, Mimi Xu, Matthew A. Lalli, John B. Hogenesch, Thomas Weimbs

**Affiliations:** Department of Molecular, Cellular, and Developmental Biology and Neuroscience Research Institute, University of California, Santa Barbara, CA, USA.; Department of Systems Pharmacology and Translational Therapeutics, Perelman School of Medicine, University of Pennsylvania, Philadelphia, PA, USA; Center for Chronobiology, Cincinnati Children’s Hospital Medical Center, Cincinnati, OH, USA

**Author notes:** Correspondence to: Thomas Weimbs, PhD, Department of Molecular, Cellular & Developmental Biology, University of California Santa Barbara, Santa Barbara, CA 93106-9625, USA. Contact: Thomas Weimbs, PhD, Department of Molecular, Cellular & Developmental Biology, And Neuroscience Research Institute, University of California Santa Barbara, Santa Barbara, California 93106-9610. Present address: Biogen GmbH, 85737 Ismaning, Germany (Christine Winterstein). Current address: Department of Cell Biology, Lerner Research Institute, Cleveland Clinic, Cleveland, Ohio, 44195 (Pallavi Bhattaram). Current address: Elena Reales: Centro de Biología Molecular Severo Ochoa, Consejo Superior de Investigaciones Científicas and Universidad Autónoma de Madrid, Cantoblanco, 28049 Madrid, Spain. Co-first author (Adrian J. Giovannone, Christine Winterstein).

## Abstract

Syntaxins - a conserved family of SNARE proteins - contain C-terminal transmembrane anchors required for their membrane fusion activity. Here we show that syntaxin 3 (Stx3) unexpectedly also functions as a nuclear regulator of gene expression. Alternative splicing leads to a soluble isoform, termed Stx3S, lacking the transmembrane anchor. Soluble Stx3S binds to the nuclear import factor RanBP5, targets to the nucleus and interacts physically and functionally with several transcription factors, including ETV4 and ATF2. Stx3S is differentially expressed in normal human tissues, during epithelial cell polarization, and in breast cancer vs. normal breast tissue. Inhibition of endogenous Stx3S expression leads to changes in the expression of cancer-associated genes and promotes cell proliferation. Similar nuclear-targeted, soluble forms of other syntaxins were identified suggesting that nuclear signaling is a conserved, novel function common among these membrane trafficking proteins.

## INTRODUCTION

The SNARE superfamily consists of several protein families that are related to each other by the presence of one or two conserved ~60 residue long SNARE motifs (Weimbs et al., 1997b; Weimbs et al., 1998). To mediate membrane fusion, membrane-associated cognate SNAREs interact with each other via their SNARE motifs to form highly stable SNARE complexes. This complex formation is thought to provide the driving force for membrane fusion. The crucial role of SNAREs in membrane fusion is well accepted and many mechanistic and regulatory details are well understood (Rothman, 2014; Sudhof, 2014; Wickner and Schekman, 2008).

Members of the syntaxin (Stx) family of SNAREs are central to all SNARE complexes and contain a C-terminal transmembrane domain, one copy of the SNARE motif, and an N-terminal three-helix bundle (Weimbs et al., 1997b; Weimbs et al., 1998). The human genome encodes at least 16 syntaxins that localize to their specific membrane domains or organelles to mediate membrane fusion reactions (Hong, 2005). For example, Stx1A and Stx1B are involved in synaptic vesicle fusion during neurotransmitter release (Sudhof, 2014) while Stx3 localizes to the apical plasma membrane domains of polarized epithelial cells (Delgrossi et al., 1997; Li et al., 2002; Low et al., 1996; Low et al., 2002), functions in polarized trafficking pathways and is essential to the correct establishment of cell polarity (Kreitzer et al., 2003; Low et al., 1998; Sharma et al., 2006; Weimbs et al., 1997a).

We report here the unexpected discovery of a novel function of Stx3 - and likely many other Stx family members - as a regulator of gene expression in the nucleus. We show that a soluble form of Stx3 is generated by alternative RNA splicing. This non-membrane anchored Stx3 isoform (termed Stx3S) interacts with the nuclear import factor RanBP5, undergoes nuclear translocation, and binds to and regulates several transcription factors including the ETS domain transcription factor ETV4 (also known as Pea3 or E1AF) and the leucine zipper family transcription factor ATF2. Both, ETV4 and ATF2 are implicated in carcinogenesis and metastasis (de Launoit et al., 2006; Gozdecka and Breitwieser, 2012). The Stx3S splice variant is endogenously expressed in human tissues *in vivo* and in several cell lines in culture. Its expression is up-regulated during polarization of human colon adenocarcinoma cells, and down-regulated in triple-negative breast cancer samples compared to normal human breast samples, suggesting a possible role in carcinogenesis. Inhibition of endogenous Stx3S expression leads to changes in cellular gene expression and promotes cell proliferation.

Our results demonstrate that this novel isoform of Stx3 can act as a transcriptional regulator. Besides Stx3, several other syntaxin family members also undergo similar alternative splicing events leading to non-membrane-anchored forms suggesting that nuclear regulatory functions are conserved among SNAREs of the syntaxin family. These results suggest the existence of previously unrecognized signaling mechanisms that may directly link membrane trafficking pathways with the regulation of gene expression.

## RESULTS

### Nuclear targeting of Stx3

To identify new Stx3 interaction partners, co-immunoprecipitation experiments were performed using C-terminally myc-epitope tagged Stx3 stably transfected in the MDCK renal epithelial cell line under the control of a doxycycline (DOX) inducible promoter (Sharma et al., 2006). In addition to the anticipated, known Stx3 binding partner munc-18b (Fig. 1A, b and b) a protein with a relative molecular mass of 110 kDa co-precipitated with Stx3 (Fig. 1A, b and a). By mass spectrometry (Fig. S1A) this protein was identified as the nuclear import factor RanBP5, also known as karyopherin β3, importin β3, or importin 5 (Deane et al., 1997). Interaction between exogenous myc-tagged Stx3 and endogenous RanBP5 was verified by reciprocal co-immunoprecipitation (Fig. 1B, 1C).

**Fig. 1.**
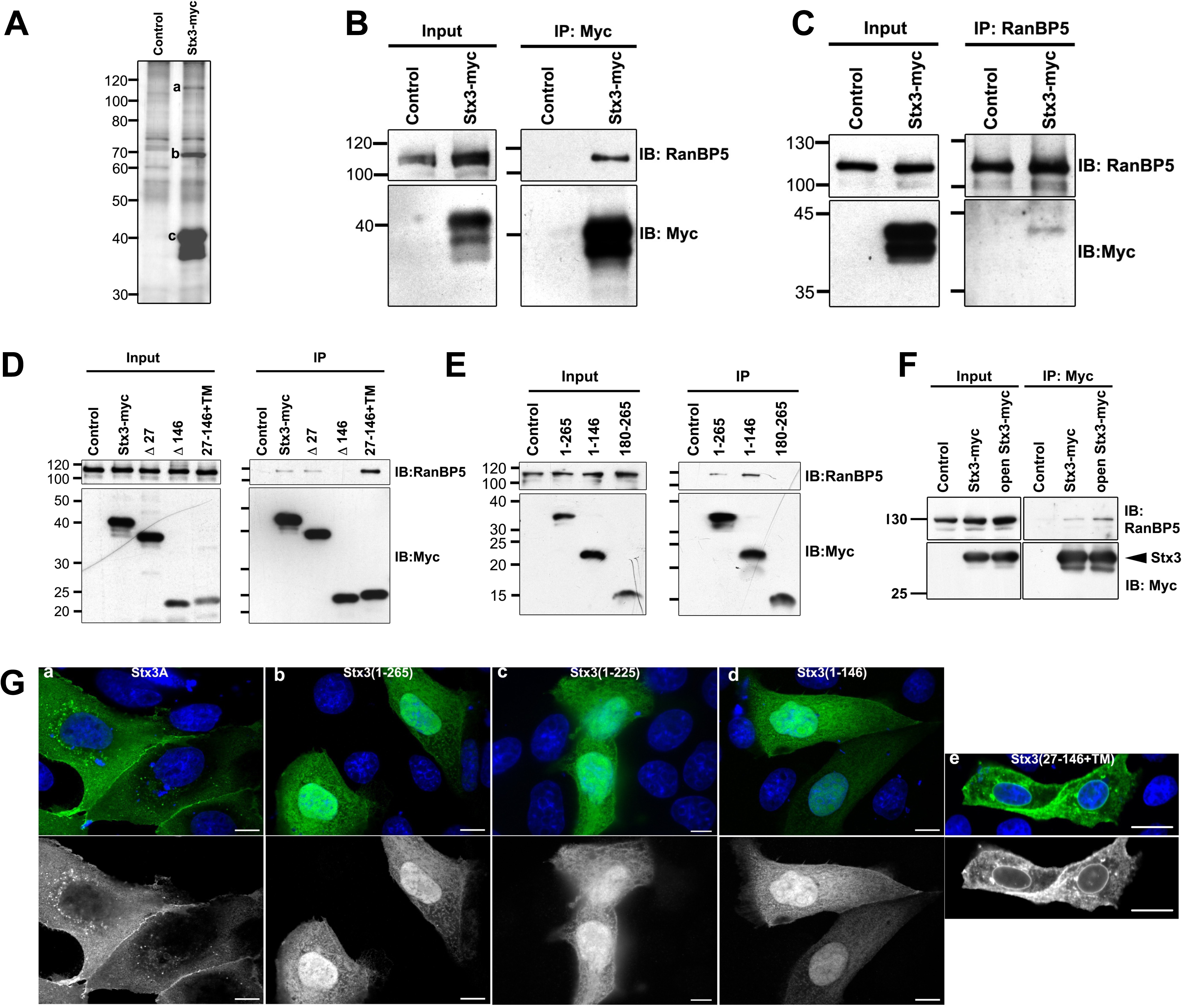
Stx3 binds to RanBP5 and targets to the nucleus. (A) Co-immunoprecipitates from MDCK lysates stably expressing myc-tagged Stx3A using an anti-myc antibody. Silver-stained bands a, b, and c were identified by MS-MS as RanBP5, Munc18b, and Stx3 respectively. (B, C) Immunoblots showing co-immunoprecipitates from MDCK lysates stably transfected with myc-tagged Stx3A and endogenous RanBP5 using an anti-myc (B) and anti-RanBP5 (C) antibody. (D, E) Immunoblots of co-immunoprecipitates from MDCK lysates transiently expressing myc-tagged Stx3A and Stx3 deletion constructs as indicated. Membrane-anchored (D) and soluble (E) Stx3 constructs were immunoprecipitated using an anti-myc antibody. (F) Lysates from HEK293T cells transiently transfected with myc-tagged Stx3A or an open mutant Stx3(L165A/E166A). IP, Immunoprecipitation. IB, Immunoblot. (G) Immunocytochemistry of MDCK cells transiently expressing the indicated myc-tagged Stx3 forms. Top row: Stx3 is stained using an anti myc antibody (green) and nuclei are stained with DAPI (blue). Bottom row: Stx3 channel only. Scale bars, 5 μm for a-d and 10 μm for e.

Using Stx3 deletion constructs (Fig. S1B), the N-terminal three-helix bundle of Stx3 was identified as the RanBP5-interacting domain (Fig. 1D, E). The apparent strength of the Stx3-RanBP5 interaction increases if the three-helix bundle of Stx3 is expressed in isolation, either in a soluble (1–146) or membrane-anchored form (27–146+TM) (Fig. 1D, E). Syntaxins are known to adopt a “closed” conformation in which the three-helix bundle folds back onto the SNARE domain to form an intramolecular helical bundle. In this conformation, syntaxins are unable to interact with other SNARE partners. In contrast, in the “open” conformation, the three-helix bundle is free and syntaxins can engage in SNARE interaction (Bock et al., 2001; Fasshauer et al., 1998; Sutton et al., 1998). To determine whether RanBP5 favors the “open” conformation of Stx3, the L165A/E166A mutation was introduced, which locks Stx3 in its “open” conformation (Dulubova et al., 1999). As shown in Fig. 1F, the L165A/E166A mutation enhances the interaction of Stx3 with RanBP5. Altogether, these results indicate that the nuclear import factor RanBP5 interacts with the three-helix bundle motif of Stx3 and that this interaction preferentially occurs when this motif is not engaged in interactions with Stx3’s SNARE motif.

We hypothesized that the interaction with RanBP5 may result in import of Stx3 into the nucleus provided that Stx3 is not membrane-anchored. To test this, membrane-anchored Stx3A and several soluble constructs lacking the membrane anchor were expressed in MDCK cells. Consistent with previously results (Li et al., 2002; Low et al., 1996; Sharma et al., 2006), membrane anchored Stx3A localizes to the plasma membrane and intracellular vesicles (Fig. 1G, a). Soluble Stx3 constructs that contain the three-helix bundle but lack the transmembrane anchor concentrate in the nucleoplasm, suggesting active nuclear import (Fig. 1G, b-d). Furthermore, a membrane-anchored mutant of Stx3 that is lacking the SNARE domain (Stx3(1-146-TM)) is prominently targeted to the nuclear envelope, in addition to other locations, but not the nucleoplasm (Fig. 1G, e). These observations suggest that Stx3 is actively transported to the nucleoplasm – presumably *via* its interaction with RanBP5 – provided it is not membrane-anchored.

### Alternative splicing leads to a Stx3 isoform lacking the trans-membrane domain

Since Stx3 is an integral membrane protein, the finding of nucleoplasmic targeting could only be of plausible significance if the membrane anchor of Stx3 was absent. Isoforms derived from alternative splicing of several syntaxins have been reported (Ibaraki et al., 1995; Martin-Martin et al., 1999), including some that lack apparent transmembrane domains (Pereira et al., 2008), but their relevance or function have remained unknown. We used BLAST to search for cDNAs of potential novel isoforms of human Stx3 in expressed sequence tag (EST) databases. Several ESTs supported the existence of an isoform lacking exon 10 (Fig. 2A). This would lead to the loss of the trans-membrane domain and expression of a truncated, soluble protein containing a unique C-terminal 12-residue peptide (Fig. 2B). We termed this novel isoform Stx3S (Stx3 “<u>S</u>oluble”). Expression of Stx3S was confirmed by RT-PCR from human embryonic kidney cells (HEK293T) (Fig. 2C), DNA sequencing of the amplified cDNA fragments, and the full-length cDNA of Stx3S was isolated and cloned. RT-PCR analysis of normal human kidney revealed that Stx3S is expressed in addition to membrane-anchored Stx3A (Fig. 2D).

**Fig. 2.**
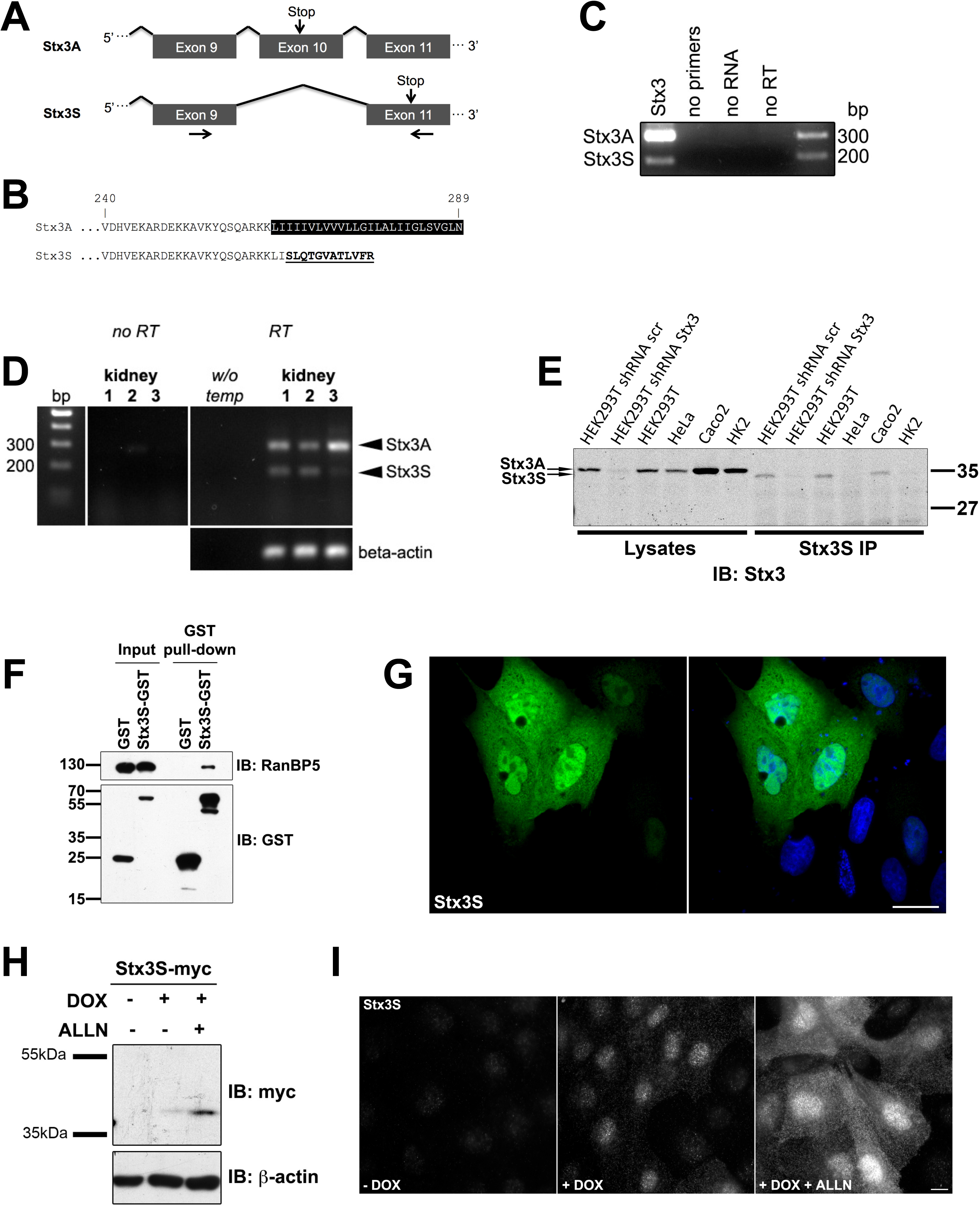
Identification of a novel splice-isoform of Stx3. (A) Schematic of exons 9 to 11 of the human Stx3 gene. Arrows mark primer pair used for RT-PCR in Fig. 2C, 2D and 3A. (B) Amino acid alignment of the C-terminal region of Stx3A and Stx3S. (C) RT-PCR of Stx3A and Stx3S in HEK293T cells. (D) RT-PCR of Stx3A and Stx3S in normal human kidney tissue from three different individuals. (E) Immunoprecipitation of Stx3S in various human cell lines including HEK293T cell line stably expressing shRNA against all Stx3 isoforms. Immunoblot using anti-Stx3 antibody shows expected size shift of Stx3S to Stx3A. (F) Co-precipitation experiment of GST-Stx3S, transiently expressed in HEK293T cells. (G) Immunocytochemistry of MDCK cells transiently expressing myc-tagged Stx3S. Stx3S is stained using an anti-myc antibody (green); nuclei are stained with DAPI (blue). Scale bar 5μm. (H) Lysates from induced or uninduced stably expressing Stx3S MDCK cells were treated with ALLN. (I) Immunocytochemistry of induced or uninduced stably expressing Stx3S MDCK cells treated with ALLN. Scale bar 5μm.

To confirm that Stx3S is expressed endogenously at the protein level, an antibody against its unique C-terminal peptide was generated and tested in a panel of human cell lines. Endogenous Stx3S protein is detectable in HEK293T and Caco2 cells (Fig. 2E) consistent with mRNA expression in these cell lines (Fig. 2C, J). Targeting an mRNA region common between Stx3A and Stx3S by shRNA diminishes expression of both Stx3 and Stx3S confirming specificity of the protein signal (Fig. 2E). At the endogenous protein level, Stx3S is expressed much lower than Stx3A and only detectable after enrichment by immuno-precipitation (Fig. 2E).

Using an *in vitro* binding assay, we verified that Stx3S readily interacts with RanBP5 (Fig. 2F) and using our expression plasmid, targets to the nucleoplasm (Fig. 2G). Treatment with the proteasome inhibitor ALLN results in significantly elevated expression levels (Fig. 2H, I) indicating that Stx3S is subject to rapid proteasomal degradation. Even at the lowest detectable expression levels, Stx3S accumulates in the nucleoplasm (Fig. 2I), indicating that its nuclear localization is not a consequence of over-expression.

Altogether, these results indicate that Stx3S is a novel isoform of Stx3 generated by alternative splicing leading to a non-membrane-anchored, nuclear-targeted protein. The Stx3S mRNA is abundantly expressed in human tissue and cell lines but the protein undergoes fast turnover compared to Stx3A.

### Regulation of Stx3S expression

To test whether expression of endogenous Stx3S may be regulated during cell differentiation, we used the colon carcinoma cell line Caco2 which are frequently used in the study of epithelial cell polarity. As these cells progress past confluence, they establish cell-cell junctions and various features of polarized morphology that are well described in the literature (Peterson and Mooseker, 1992; Rodriguez-Boulan and Nelson, 1989). Expression levels of Stx3A vs. Stx3S were monitored at different time points during the establishment of polarized epithelial morphology. In contrast to Stx3A, which does not show a difference in expression during polarization, Stx3S mRNA is upregulated after day two and maintains its expression level thereafter (Fig. 3A). At the protein level, the upregulation of Stx3S is even more pronounced with Stx3S protein detectable at day 6 but not at day 3 (Fig. 3B).

**Fig. 3.**
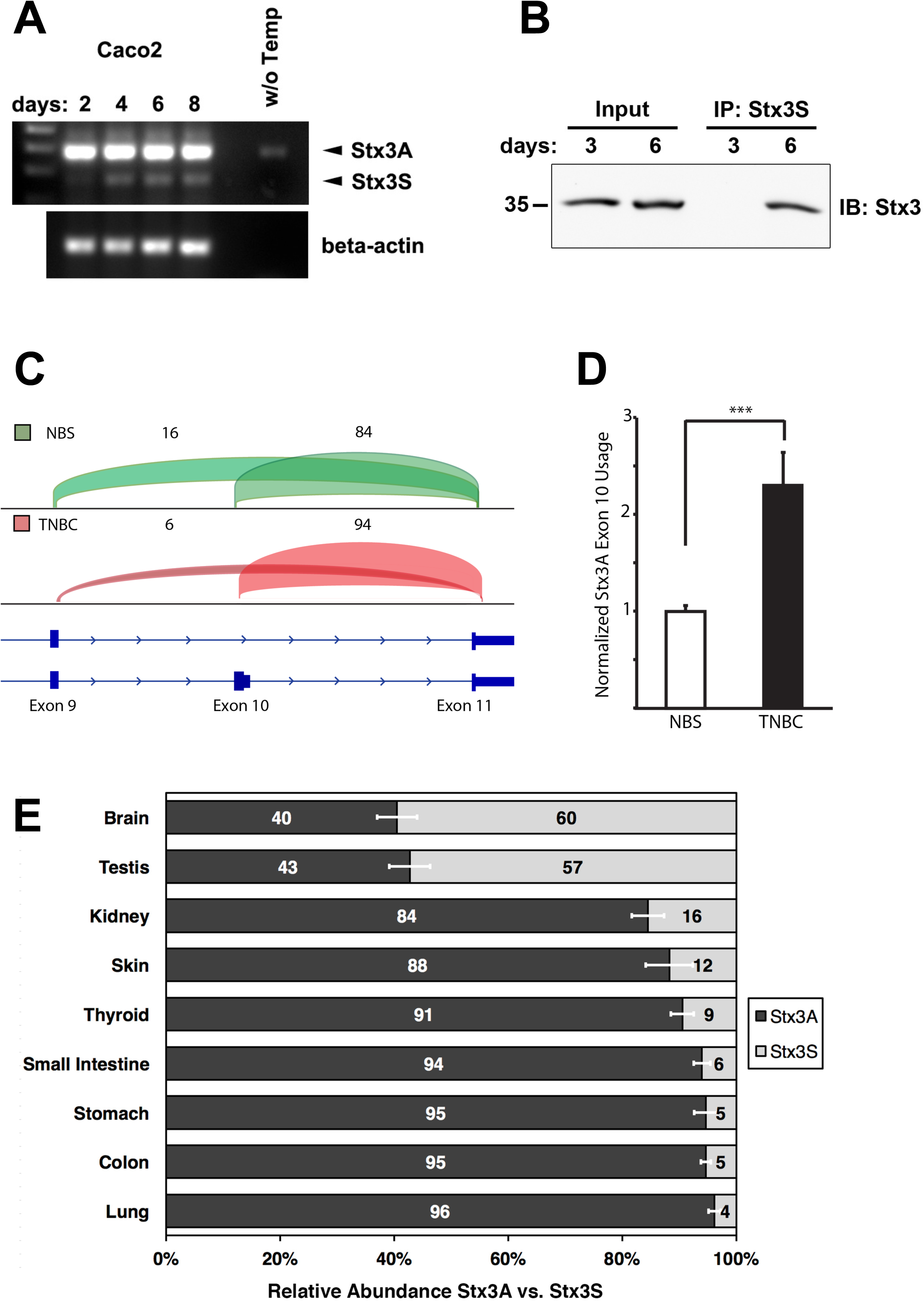
Regulation of Stx3S expression. (A) Caco2 cells were cultured for the indicated durations and analyzed by RT-PCR for Stx3A/Stx3S mRNA expression using the primer pair shown in Fig. 2A. The Stx3S transcript is notably upregulated during the transition from a sub-confluent, highly proliferative culture (day 2) to confluent and post-confluent cultures (later days). (B) Immunoprecipitation with Stx3S-specific antibody from Caco2 cells cultured for indicated days. Immunoblot used total Stx3 antibody. The IP lanes are 20x enriched over the input lanes. Therefore, only the more abundant Stx3A is visible in the input lanes. (C) Genome browser view of transcriptome analysis of Stx3 transcripts in triple-negative breast cancer (TNBC) compared to normal breast samples (NBS). n_NBS_ = 3, n_TNBC_ = 6. Transcripts containing the exon 10/11 junction are unique to Stx3A whereas transcripts containing the exon 9/11 junction are unique to Stx3S. The percentages of reads mapping to junctions 9/11 and 10/11, respectively, are indicated by numbers and visualized by brackets of heights proportional to their abundance. Stx3S is notably downregulated, compared to Stx3A, in TNBC. (D) *** *P*<0.001. Data are shown as mean ±SEM. Statistical analysis was performed using *DEXSeq* package. (E) Relative Stx3A vs. Stx3S isoform expression based on analysis of splice junction usage from RNA sequencing data of multiple human tissues. Data are shown as the mean ± SEM.

To investigate whether alternative splicing between Stx3A and Stx3S may also be regulated *in vivo* and may play a role in disease, the expression profiles of Stx3 were examined in transcriptomes from human normal breast samples (NBS) and triple-negative breast cancer (TNBC) (Eswaran et al., 2013; Horvath et al., 2013). Examination of RNA-seq reads aligning to splice junctions specific to Stx3A (exon 10/11 junction) and Stx3S (exon 9/11 junction) transcripts, respectively, revealed a notable depletion of Stx3S in TNBC relative to normal samples (Fig. 3C). A relative increase in the usage of the Stx3A-specific exon 10 was detected in TNBC samples, indicating a corresponding downregulation of Stx3S (Fig. 3D). This indicates that expression of Stx3S is downregulated in TNBC.

Examining transcriptomes of various normal human tissues also revealed striking differences in the alternative mRNA splicing leading to Stx3S vs. Stx3A transcripts (Fig. 3E). Tissues with the largest relative expression of Stx3S are testis and brain with over 50 percent of the Stx3 transcript population. This is in contrast to other tissues, e.g. lung, which exhibits a drastically lower abundance of Stx3S mRNA at 4 percent.

Altogether, these results indicate that Stx3S is a widely expressed isoform whose expression is controlled by tissue-specific regulation of alternative RNA splicing, and is downregulated in tumor cells *in vivo* and in rapidly proliferating carcinoma cells *in vitro*.

### Soluble Stx3 interacts physically and functionally with transcription factors

The discovery of a soluble, nuclear-targeted, rapidly degraded isoform of Stx3 led us to hypothesize that Stx3S may function as a signaling protein that may regulate gene expression. To test this possibility, a classic Gal4 based transcriptional transactivation assay was employed (Vecchi et al., 2001). Constructs encoding the Gal4 DNA-binding domain fused to membrane-anchored Stx3A or cytoplasmic Stx3(1–265), respectively, were co-expressed with a reporter plasmid encoding the luciferase gene under control of a Gal4-responsive promoter. Expression of Gal4-Stx3(1–265) resulted in fourfold transactivation over Gal4 alone or the membrane-anchored Gal4-Stx3A construct (Fig. 4A). This finding suggests that the cytoplasmic Stx3 domain either has intrinsic transcriptional activation capability and/or interacts with other transcription factors.

**Fig. 4.**
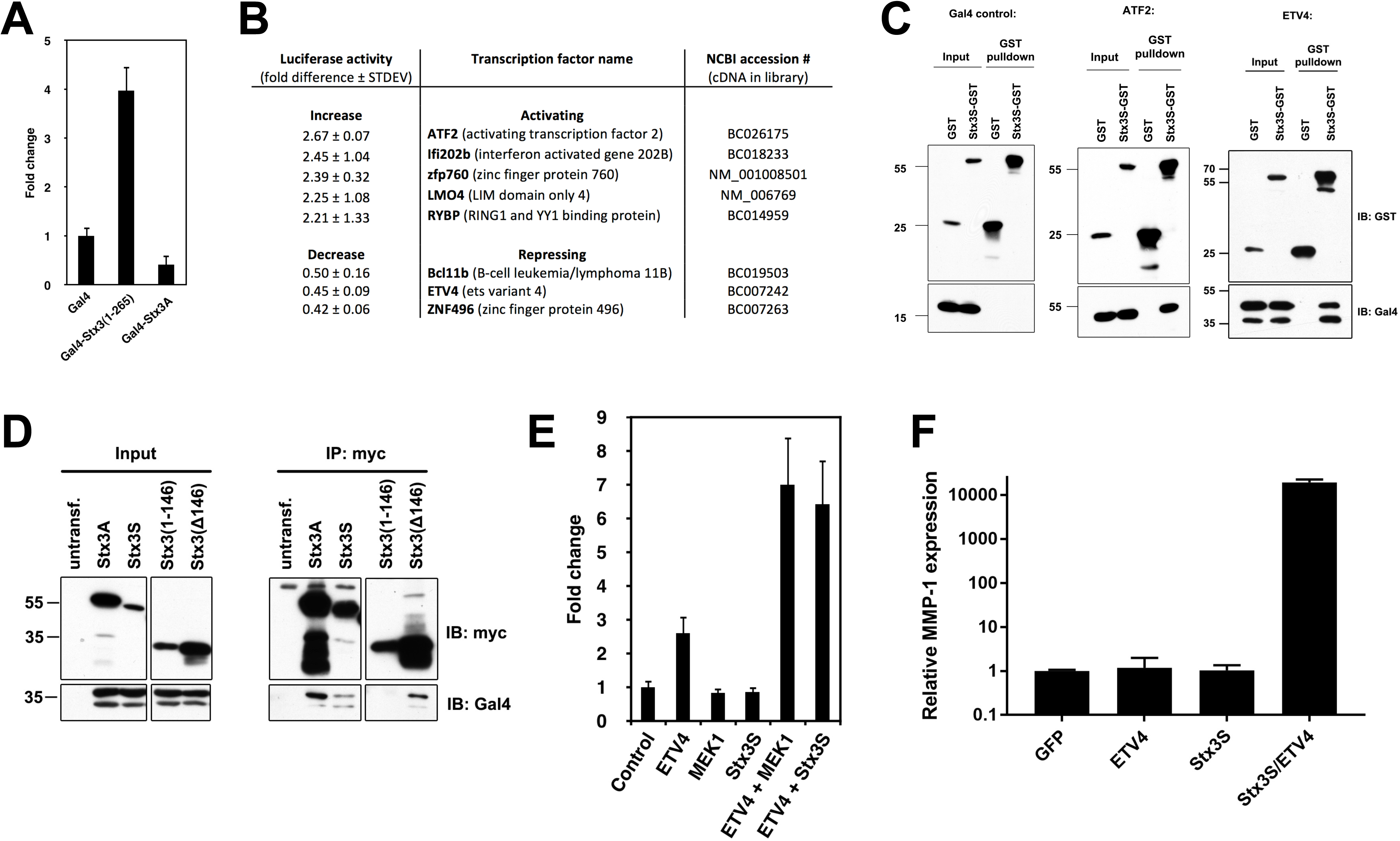
Stx3S acts as a transcriptional co-activator of ETV4. (A) COS-7 cells were transiently transfected with soluble Stx3 (1–265) or membrane anchored Stx3A fused to the DNA binding domain of Gal4. Transcriptional activity was measured using a luciferase reporter construct containing Gal4 binding elements (n=3). (B) Table showing transcription factors whose activity was most strongly altered by Stx3S in the co-activator trap screen. (C) GST-Stx3S and Gal4-ETV4 or Gal4-ATF2 were co-expressed in HEK293T cells and co-precipitated from lysates using glutathione-Sepharose. (D) Myc-tagged Stx3A, Stx3S, and Stx3 deletion constructs were co-expressed with Gal4-ETV4 in HEK293T cells. Cells were harvested for immunoprecipitation with anti-myc and blotted for Gal4 or myc, as indicated. (E) ETV4-reporter assay in HeLa cells, showing increased luciferase activity after co-transfection of ETV4 with constitutively active MEK1 or Stx3S (n=3). A non-responsive β-galactosidase plasmid was used as an internal control and signals were normalized to β-galactosidase. (F) HEK293T cells were transiently transfected with GFP (control), ETV4, Stx3S, or Stx3S/ETV4, stimulated with PMA for 4h, and harvested for reverse transcriptase and qPCR to quantify endogenous MMP-1 expression. MMP-1 expression was normalized to GAPDH. Bars represent the means and SEM of triplicate samples. Data are representative of three independent experiments.

To identify transcription factors that may functionally interact with Stx3S, a high-throughput, cell-based screening assay, termed "co-activator trap” was employed (Amelio et al., 2007). An arrayed library of 1465 sequenced-verified plasmids coding for mouse and human transcription-related proteins fused to the Gal4 DNA binding domain was co-expressed with a Gal4 UAS::luciferase reporter in the presence or absence of Stx3S. Several transcription factors were identified whose activity was affected by Stx3S. The top eight that exhibited the strongest co-activation or co-repression by Stx3S were selected for validation and further analysis (Fig. 4B). To confirm binding interactions, a GST-Stx3S fusion protein was expressed together with each of these transcription factors (as fusion proteins with the Gal4 DNA binding domain). Six out of these eight transcription factors (ETV4, ATF2, Ifi202b, Znf496 and LMO4) co-precipitated with GST-Stx3S indicating that they form stable complexes (Fig. 4C, S2). ETV4 and ATF2 showed the most robust interactions.

### Stx3S co-activates ETV4

ETV4 was selected for further analysis because its interaction with Stx3S was very consistent and because both ETV4 and Stx3 have been shown to play roles in renal epithelial differentiation. ETV4 is essential for branching morphogenesis during renal development (Kuure et al., 2010; Lu et al., 2009) and associated with several epithelial cancers (de Launoit et al., 2006). Stx3A is highly expressed in renal tubule epithelial cells (Li et al., 2002) and required for the establishment of epithelial cell polarity (Sharma et al., 2006).

Immunoprecipitation experiments with truncated Stx3 constructs revealed that the C-terminal half of Stx3 interacts with ETV4 (Fig. 4D) suggesting that the N-terminal three-helix bundle may remain available for binding to RanBP5.

To test whether Stx3S acts on ETV4-regulated promoters, a luciferase reporter gene controlled by five ETV4 binding elements was used. Constitutively active MEK1 served as a positive control, since ETV4 activity is enhanced in response to the ERK/MAP kinase signaling pathway (O'Hagan et al., 1996). Expression of either, constitutively active MEK1 or Stx3S caused a significant increase in ETV4-dependent reporter gene activity (Fig. 4E). Next, we examined whether Stx3S can regulate the expression of a native, endogenous ETV4 target gene. ETV4 has previously been shown to increase MMP-1 expression in epithelial cancers (Bosc et al., 2001; Keld et al., 2010). Expression of Stx3S together with ETV4 in HEK293T cells leads to extremely strong endogenous MMP-1 expression compared to either Stx3S or ETV4 alone (Fig. 4F). Altogether, these results demonstrate that Stx3S physically interacts with ETV4 and acts as a co-activator of ETV4-regulated transcription.

### Loss of endogenous Stx3S leads to changes in gene expression and increased cell proliferation

In order to demonstrate a function of endogenous Stx3S, independent of its overexpression, we specifically silenced expression of Stx3S without affecting Stx3A. To achieve this, multiple shRNA sequences flanking the unique exon 9/11 border were tested in HEK293T cells. shRNA sequence #C4 led to efficient and specific knockdown of Stx3S expression and was used for all subsequent experiments (Fig. S3A, S3B). Stable Caco2 and HEK293T cell lines were generated with the Stx3S-specific shRNA or scrambled control shRNA (Fig. 5A). To identify Stx3S-regulated target genes in a non-biased fashion the gene expression profiles between Caco2 cells with silenced Stx3S expression and control Caco2 cells were compared. Numerous genes were found to be differentially regulated in the absence of endogenous Stx3S and those genes with the largest difference are shown in Fig. 5B. Many of these affected genes are predicted to be regulated by ATF2 (Kent et al., 2002) which we had found to physically and functionally interact with Stx3S (Fig. 4B, C). The Stx3S-dependent regulation of three of the genes predicted to be influenced by ATF2 (FNTA, KDM2B, and SART3) were confirmed with qPCR (Fig. 5C). These results indicate that loss of endogenous Stx3S leads to distinct changes in gene expression consistent with a role as a regulator of transcription factors including ATF2.

**Fig. 5.**
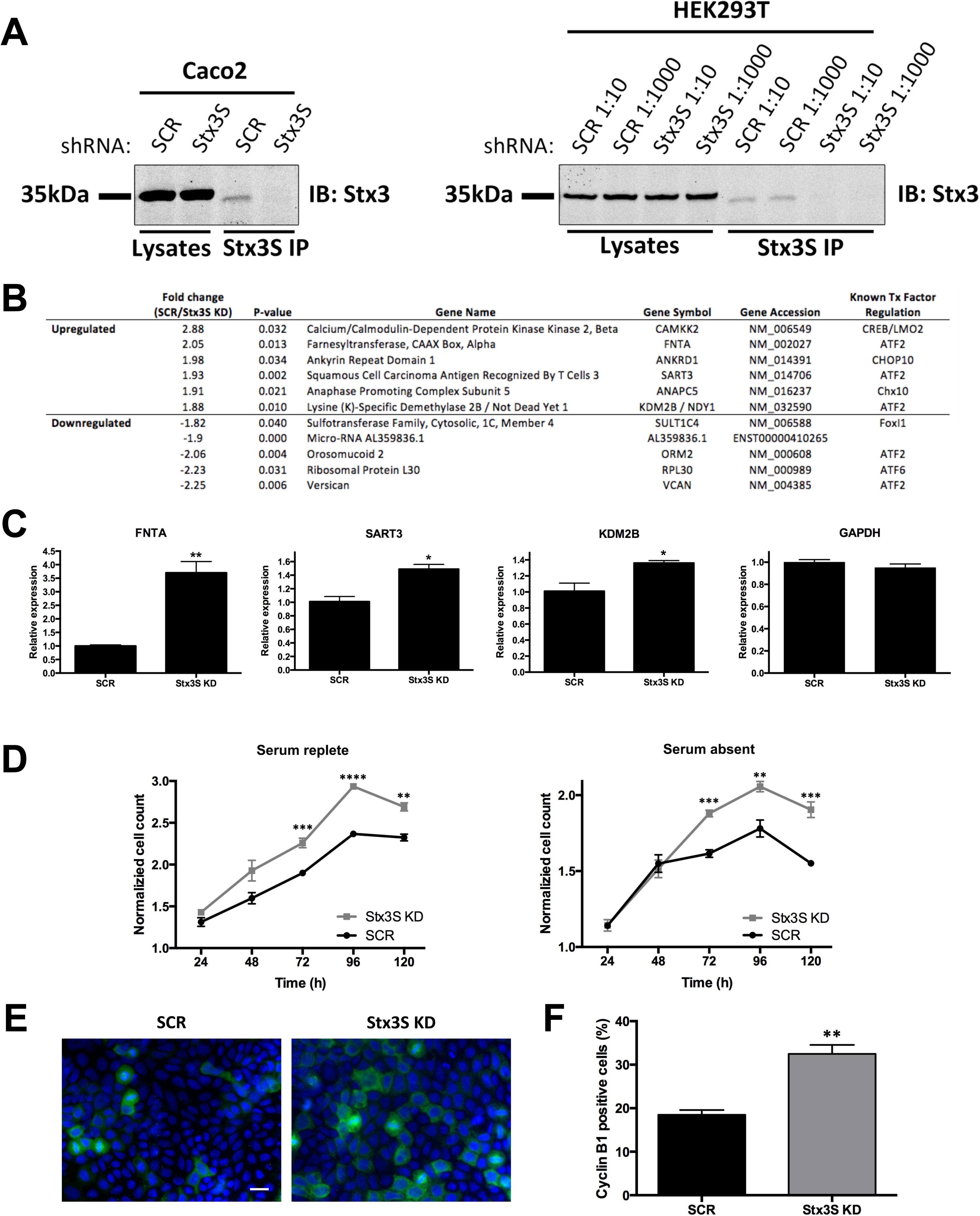
Stx3S acts as a transcriptional co-activator of ATF2 and affects cell proliferation. (A) Immunoprecipitation of Stx3S in Caco2 and HEK293T cells, respectively, stably transduced with lentivirus delivering shRNA #C4 targeting Stx3S (see Fig. S3B). HEK293T cells were transduced with two different dilutions of viral media leading to similar knock-down efficiencies. (B) RNA from Stx3S KD Caco2 cells was subjected to Affymetrix microarray analysis. Statistically significant transcripts with the largest fold-difference compared to control cells are shown in the table, including the transcription factors predicted to associate with them (as projected by SABiosciences' Text Mining Application and the UCSC Genome Browser)(Kent et al., 2002). (C) qPCR of select genes from microarray analysis. All genes normalized to β-actin. Bars represent the means and SEM of triplicate samples. *P< 0.05; **P<0.01 (Student’s unpaired *t*-test) Data is representative of three independent experiments. (D) Growth curve of Stx3S KD or shRNA scrambled control (SCR) Caco2 cells determined by CellTiter-Glo cell viability assay, normalized to 24 hour reading. Growth curves were determined in the presence or absence of serum side-by-side. Data points represent the means and SEM of quadruplicate samples. **P<0.01; ***P<0.001; ****P<0.0001 (Student’s unpaired *t*-test). (E) Representative immunocytochemistry images of 5-day cultures of SCR or Stx3S KD Caco2 cells stained for Cyclin B1 (green) and DAPI (blue). Scale bar 20μm. (F) Cyclin B1 positive proportion for SCR and Stx3S KD cells. Bars represent the means and SEM of triplicate samples. **P<0.01 (Student’s unpaired *t*-test)

We assessed whether loss of endogenous Stx3S expression may lead to distinct cellular phenotypes. As shown in Fig. 5D, knock-down of Stx3S expression leads to an increase in cell proliferation in Caco2 cells both in media containing serum and in serum-free conditions. Increased proliferation is validated by a 75% increase of cyclin B1 (M phase) positive staining observed in 5-day cultures (Fig. 5E, F). No difference in apoptosis is seen by cleaved caspase-3 staining (Fig. S4). These results, together with the finding that Stx3S expression is downregulated in rapidly proliferating Caco2 cells (Fig. 3A, B) and downregulated in breast cancer tissue (Fig. 3C, D) suggests that Stx3S may play a role in dampening proliferation of epithelial cells and may therefore be a tumor suppressor.

### Soluble splice variants of other syntaxin genes

Evidence for a handful of syntaxin variants lacking transmembrane anchors has previously been reported (Jagadish et al., 1997; Nakayama et al., 2012; Nakayama et al., 2003; Nakayama et al., 2004; Pereira et al., 2008; Quinones et al., 1999), but their functions are unknown. To understand whether a conserved mechanism may be involved in the generation of these alternative transcripts from different syntaxin genes, we analyzed the mechanisms leading to the previously reported “soluble” variants of Stx1A, Stx1B, and Stx2. Furthermore, human EST databases were searched for possible evidence of novel “soluble” variants of other syntaxin genes. In addition to Stx3S reported here, we identified alternative transcripts - supported by numerous ESTs - leading to novel variants of other syntaxins lacking C-terminal transmembrane anchors, including Stx4 and Stx10 (Fig. 6A). Because the nomenclature of syntaxins is frequently inconsistent we propose to name any syntaxin variant lacking a C-terminal transmembrane anchor by appending the letter S (soluble) to the original gene name as shown in Fig. 6A.

**Fig. 6.**
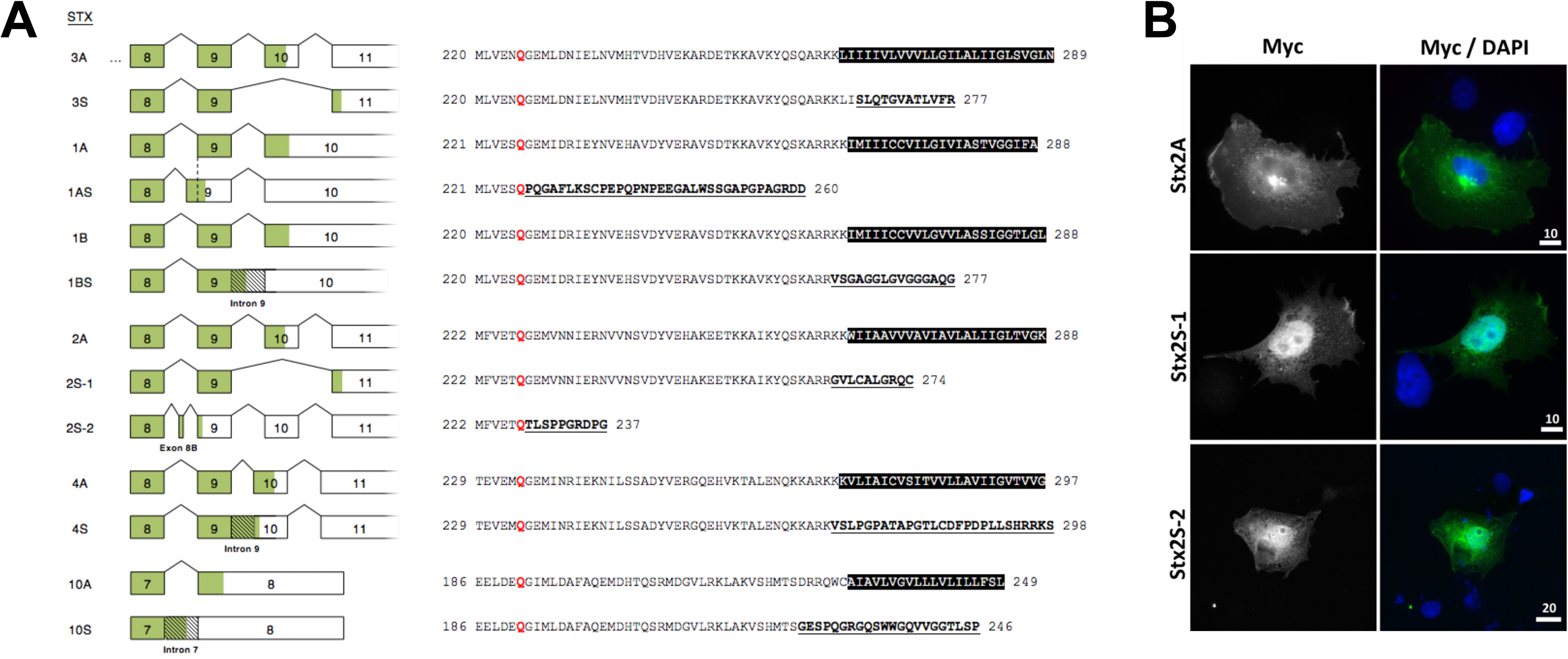
Soluble, nuclear-targeted isoforms are common among the syntaxin gene family. (A) Schematic of previously or newly identified splice variants of various syntaxin genes coding for soluble proteins lacking C-terminal transmembrane anchors. For consistency, “soluble” isoforms are termed here by appending the letter “S”. RNA splicing patterns for the canonical membrane-anchored variants are compared to splicing for the “S” variants. Boxes represent exons with coding regions shaded in green. C-terminal protein sequences are shown with transmembrane domains highlighted in black, and the novel sequences of “S” isoforms shown underlined and bolded. The conserved glutamine residue between all syntaxin family members is highlighted in red. Soluble isoforms human Stx1AS (originally termed Stx1C) (Jagadish et al., 1997), human Stx1BS (originally termed Stx1B-ΔTMD) (Pereira et al., 2008), rat Stx2S-1 (originally termed Stx2C)(Quinones et al., 1999), rat Stx2S-2 (originally termed Stx2D) (Quinones et al., 1999) have been described previously. Soluble isoforms human Stx3S, human Stx4S, human Stx10S were identified in this study. (B) Immunocytochemistry of COS-7 cells transiently expressing myc-tagged Stx2A, Stx2S-1 or Stx2S-2, respectively. Cells were stained with anti-myc antibody (green) and DAPI (blue). Scale bars in μm.

The alternative transcripts coding for these soluble syntaxins are generated by exon skipping (Stx3S, Stx2S-1), use of an alternative 3′ acceptor site (Stx1AS), use of a cryptic exon (Stx2S-2), or intron retention (Stx1BS, Stx4S, Stx10S), respectively (Fig. 6A). In all cases, this leads to the replacement of the transmembrane domain with short, hydrophilic sequences that exhibit no apparent sequence similarity among them.

While Stx1BS has been reported (Pereira et al., 2008) to exhibit nuclear localization similar to Stx3S, no targeting information is available for the other isoforms. To investigate whether nuclear targeting may be a common feature of soluble syntaxin isoforms, Stx2S-1 and Stx2S-2 were investigated. Both isoforms target to the nucleoplasm when expressed *in vitro* (Fig. 6B). Altogether, these results suggest that mammalian syntaxin genes are commonly expressed as alternative transcripts leading to soluble isoforms that undergo nuclear targeting.

## DISCUSSION

Here we demonstrate that a novel isoform of Stx3 acts as a transcriptional regulator in the nucleus. This function requires the lack of the C-terminal trans-membrane anchor due to alternative splicing to generate the soluble Stx3S. This is an unexpected, novel function for a member of the SNARE family. However, it appears likely that nuclear functions may not be unique to Stx3 but represent a characteristic that is common among many members of this SNARE family. Our finding that “soluble” splice variants of numerous other syntaxins exist supports this. A splice-isoform of human Stx1B (termed Stx1BS in Fig. 6A) was reported to localize to the nucleus of various tumor and non-tumor cell-types (Pereira et al., 2008). Stx17 has been reported to localize to the nucleus by immunohistochemistry in several cell types (Zhang, 2005) but it is currently unknown whether this may be due to the existence of an alternatively spliced variant. A human Stx1A splice variant (termed Stx1AS in Fig. 6A) is expressed in numerous tissues and cell types (Jagadish et al., 1997; Nakayama et al., 2003) but its subcellular targeting has not been thoroughly investigated. We demonstrate that Stx2S-1 and Stx2S-2, two previously reported “soluble” variants, exhibit nuclear localization (Fig. 6B). In addition, we identified novel variants of Stx4 and Stx10 that lack transmembrane domains (Fig. 6A). While most of the identified soluble syntaxin variants diverge from the membrane-anchored variants immediately adjacent to the start of the transmembrane domain, two isoforms (Stx1AS and Stx2S-2) instead diverge immediately after a highly conserved glutamine in the middle of the SNARE motif (Fig. 6A). In the case of Stx2, both types of soluble variants exist and both exhibit nuclear targeting (Fig. 6B). The C-terminal, hydrophilic peptide extensions of the soluble Stx splice isoforms (Fig. 6A) do not exhibit any apparent homology to each other or to other known motifs, and are not required for nuclear targeting. It remains to be elucidated whether these short sequences have any function besides replacing the trans-membrane anchors.

We cannot formally rule out that soluble Stx3 may also have a SNARE-related function in regulating membrane fusion, e.g. by acting as a dominant-negative inhibitor of SNARE interactions. However, experimental experience indicates that recombinant, soluble versions of SNAREs only act as dominant-negative inhibitors at very high, artificial expression levels (Low et al., 2003a; Weimbs et al., 2003). We consider it unlikely that the extremely low, endogenous expression level of Stx3S described here would affect normal SNARE function, especially given that Stx3S is targeted to the nucleus and is therefore separated from cytoplasmic SNAREs.

We found that the expression of Stx3S is up-regulated in Caco2 colon carcinoma cells as they transition from a rapidly proliferating phenotype to a fully confluent, polarized epithelial phenotype (Fig. 3A, B). Similarly, the expression of Stx3S is higher in normal breast epithelium compared to triple-negative breast cancer (TNBC) samples (Fig. 3C, D). TNBC, a particularly aggressive and therapy-resistant form of breast cancer, is characterized by elevated expression of farnesyltransferase alpha (FNTA) (Singha et al., 2013) and the histone demethylase KDM2B/NDY1 (Kottakis et al., 2014), two genes whose expression we have found to increase when Stx3S is silenced (Fig. 5B, C). This connection is further supported with the observed increase in proliferation of Caco2 cells lacking Stx3S (Fig. 5D-F). Altogether, these results suggest that Stx3S may regulate epithelial phenotypes important in carcinogenesis. A role in tumorigenesis has also been proposed for Stx1BS (Pereira et al., 2008) which exhibits nuclear localization in human glial tumors and its expression correlates with a worse outcome in brain tumor patients.

The low protein expression level of Stx3S observed appears to be due to rapid proteasomal degradation (Fig. 2H, I). Such rapid turnover would be typical for signaling proteins, and may be the reason why the nuclear localization of Stx3 and other syntaxins has previously escaped detection. Altogether, our results have uncovered a novel and unforeseen signaling pathway. Syntaxins are a conserved family of SNARE proteins involved in every intracellular membrane trafficking step in eukaryotic cells. Signaling by soluble syntaxins may be of fundamental importance to a wide variety of cellular functions.

## EXPERIMENTAL PROCEDURES

### Cell culture and transfections

MDCK cells were cultured as described previously (Low et al., 2000). HEK293T, Caco2, and COS-7 cells were cultured in DMEM with 10% FBS, penicillin, and streptomycin. TurboFect (Thermo Scientific) or Fugene 6 (Promega) were used for transient transfections. Calcium phosphate precipitation was used for stable transfection of MDCK cells. Stable cell-lines were generated as previously described (Sharma et al., 2006). Caco2 and HEK293T cells stably expressing shRNA against Stx3S were generated by transduction with lentiviral vectors followed by selection with puromycin.

### Plasmids and mutagenesis

All Stx3 constructs are based on human Stx3 as described (Sharma et al., 2006). cDNAs of full-length or truncated syntaxins were inserted into pcDNA4/TO/2xmyc/His as previously described (Low et al., 2003b). Point mutants of Stx3 were generated by site directed mutagenesis using primers containing the mutation. GFP-Stx3 was constructed by insertion of the cDNA of Stx3 into pEGFP-C1 (Clontech). GST fusion proteins for mammalian expression were constructed by insertion of GST into pcDNA4/TO/myc/His, followed by subcloning of Stx3S upstream of GST. pCMX-Gal4(DBD) encodes Gal4 DBD (amino acids 1–147) (Perlmann and Jansson, 1995). pCMX-Gal4(DBD)-wtStx3 and pCMX-Gal4(DBD)-Stx3(1–265) were constructed by inserting Stx3A or Stx3(1–265) into pCMX-Gal4(DBD). Gal4(DBD) fused to various transcription factors have been described (Amelio et al., 2007). Mouse pRSV-Pea3, FLAG-Pea3, and 5xPea3-LUC were kindly provided by John Hassell (McMaster University, Hamilton, Ontario). Human pFLAG-E1AF coding for ETV4 was kindly provided by Andrew Shamrocks, (University of Manchester, UK) has been described previously (Nishida et al., 2007). MEK1 encodes constitutively active MEK-1(ΔNS218E-S222D) (kindly provided by Dr. Roger Davis, University of Massachusetts, Worcester, MA). All constructs were verified by sequencing. Stx2 in pcDNA4 backbone was made previously (Low et al., 2003b).

### shRNA and lentivirus

Candidate shRNA sequences (Fig. S3A) against Stx3S were designed to have restriction sites appropriate for integration into the pGreenPuro plasmid from System Biosciences. Ligation was confirmed by restriction digestion. pGreenPuro shRNA plasmids were used with pMDG envelope and p8.91 packaging plasmids in HEK293T cells to generate lentiviral particles.

### Antibodies and Immunoprecipitation

Hybridoma cells for the 9E10 anti-myc epitope tag antibody were obtained from the American Type Culture Collection (ATCC, Manassas, VA). Anti-β-actin, AC-15 (Sigma), used for immunoprecipitations and western-blotting; anti-Myc, 4A6 (Millipore) and anti-Cyclin B1, D5C10 (Cell Signaling) were used for immunofluorescence; anti-GFP, JL-8 (BD Bioscience); anti-GST Gt pAb (GE Healthcare); anti-Gal4 DBD, sc-510 (Santa Cruz); anti-RanBP5, sc-11369 (Santa Cruz); anti-Pea3, sc-113 (Santa Cruz); a mouse monoclonal antibody was developed against the cytoplasmic domain of rat Stx3 (clone 1–146, licensed to EMD Millipore). Secondary antibodies conjugated to DyLight 488 or 594 and HRP were obtained from Thermo Scientific and Jackson ImmunoResearch Laboratories, respectively. Rabbit anti-Stx3S anti-serum was developed against a peptide specific to the C-terminus of Stx3S.

For RanBP5 co-immunoprecipitations, MDCK cells expressing Stx3A or COS-7 cells transiently transfected with truncated Stx3 constructs were lysed in buffer containing 50mM Hepes, pH 7.4, 150mM potassium acetate, and 0.5% Triton X-100. Myc-tagged Stx3 was immunoprecipitated by using anti-myc antibody 9E10 crosslinked to Protein G Sepharose. For GST-pulldowns and co-immunoprecipitations of Gal4-fusion proteins from the co-activator trap screen, HEK293T cells were transiently transfected using Turbofect and plasmids of interest. Cells were lysed in buffer containing 20mM Tris, 1% Triton X-100, 150mM NaCl, protease, and phosphatase inhibitors. Cells were sonicated for 15 sec on ice using a probe sonicator at 20% output, 40% duty cycle. Stx3S-GST and myc-tagged Stx3 constructs were precipitated with Glutathione conjugated beads and anti-myc (9E10) respectively. Precipitated proteins were eluted in 2xSDS sample buffer and analyzed by SDS-PAGE and western-blotting. For endogenous Stx3S immunoprecipitations, cells were lysed in buffer containing 25mM Hepes, pH 7.4 (KOH buffered), 125mM KOAc, 2.5mM MgOAc, 5mM EGTA, 1mM DTT, 0.5% Triton X-100, and protease inhibitors. Lysates were needled and centrifuged at 17,900g. Supernatants were pre-cleared with CL2B sepharose (GE Healthcare) and incubated with rabbit anti-Stx3S serum overnight at 4C. Antibodies were precipitated with Protein A Sepharose, eluted in 2xSDS sample buffer, subjected to SDS-PAGE and western-blotting.

### Mass spectrometry

C-terminally myc-tagged Stx3A was expressed in stably transfected, DOX-inducible MDCK cells, stably transfected MDCK cells cultured on 15 cm dishes, and uninduced cells served as a negative control. Proteins were immunoprecipitated using anti-myc antibody (clone 9E10) covalently linked to protein G Sepharose using dimethyl pimelimidate. Eluted proteins were analyzed by SDS-PAGE and silver staining. Identification of proteins by mass spectrometry was done as previously described (see Supporting Information in (Talbot et al., 2011). Briefly, excised bands were washed with 50% ethanol / 5% acetic acid containing DTT followed by alkylation with iodoacetamide. Samples were digested with trypsin and analyzed by LC-tandem mass spectrometry. Mass and CID spectra were gathered, compared to the NCBI nonredundant database, and confirmed manually.

### Immunocytochemistry

MDCK or Caco2 cells were fixed in 4% paraformaldehyde for 15 min. After quenching in PBS containing 75 mM ammonium chloride and 25 mM glycine, cells were permeabilized with PBS containing 3% BSA and 0.2% Triton X-100. Cells were incubated with primary antibodies overnight at 4C, followed by fluorescence-labeled (DyLight 488) secondary antibodies for 1 h at 37ºC. Images were acquired at room temperature using an Olympus IX81 microscope, a UplanSApo 60x/1.35 lens, Immersol 518 F oil (Zeiss), a Retiga EXi camera (QImaging) and IPLab software (BD Biosciences). Images were processed using Adobe Photoshop CS5 for linear histogram adjustments, and to assemble composite figures. COS-7 cells were transiently transfected with pcDNA4-Stx2A-myc, pcDNA4-Stx2S1-myc, or pcDNA4-Stx2S2 plasmids and processed as above for immunocytochemistry.

### RNAseq data analysis

RNA sequencing data derived from human primary breast tissue or tumor samples were downloaded from the Gene Expression Omnibus (accession GSE52194) and aligned to the human genome (hg19) using default settings of TopHat (v2.0.10). As these data were generated specifically to examine differential splicing in cancer, we examined the splice pattern of Stx3 in these data (n_NBS_ = 3, n_TNBC_ = 6). Stx3 exons were counted using *htseq* and differential exon usage was determined with the *DEXSeq* package in R. Splice reads mapping to exon 11 were filtered from TopHat’s junction.bed houtput, merged by sample type (tumor vs. normal) and visualized using the Integrative Genomics Viewer (IGV). As spliced reads mapping to exon 11 align either to the junction between exon 9 and 11, or between exon10 and 11, these junction counts represent the abundance of exon 10 exclusion and inclusion isoforms respectively.

RNA sequencing data from 95 human individuals representing 27 different tissues was obtained from the European Nucleotide Archive (Study: ERP003613). Tissues of interest were imported into Illumina’s BaseSpace cloud server using the SRA Import application and aligned to the human genome (hg19, RefSeq) using the TopHat Alignment application, v1.0.0. Splice junctions between exons were visualized and quantified using the Integrative Genomics Viewer (IGV) Sashimi Plot, version 2.3.68. The average number of spliced reads that aligned to the junctions between exon 9–10 and 10–11 represent the abundance of Stx3A. Spliced reads that aligned to the junction between exons 9 to 11 represent Stx3S.

### Luciferase assays

Gal4 transactivation assays were performed as described (Vecchi et al., 2001). Briefly, COS-7 cells were transiently transfected in triplicate with the different Gal4-fusion constructs, Gal4-LUC reporter, and β-Galactosidase plasmid as a transfection control. Cells were lysed in luciferase lysis buffer (Promega) 48h after transfection. Luciferase activity was measured on identical amounts of total cellular lysates using a commercial kit (Promega) and normalized to β-Galactosidase activity. In addition, lysates were analyzed by western-blotting with anti-Gal4 antibodies to verify the levels of expression of the various proteins.

A cDNA library of ~1400 Gal4-tagged transcription factors (Amelio et al., 2007) was screened with either empty vector or full length Stx3S in HEK293T cells. The transcription factor library was purified from glycerol stocks with the NucleoSpin 96 Plasmid DNA prep kit (Macherey Nagel) and spotted into 384-well plates. Library screening was done in reverse transfection format by adding serum-free DMEM containing Fugene 6 (Promega), pG5-luc (Promega), and either pcDNA4 or Stx3S to each well. Following a 20–30 minute incubation, of DMEM/20% FBS containing 10,000 HEK293T cells was added to each well. Cells were cultured for 24h and luciferase levels were measured using Britelite Plus (Perkin Elmer) on an Analyst HT plate reader (Molecular Devices).

For ETV4 reporter assays, HEK293T or HeLa cells were plated on 96-well plates. 16h later, cells were transfected with 5xPea3 RE-LUC, ETV4, Stx3 and/or MEK1 construct and β-Gal per well. One day after transfection, cells were lysed in passive lysis buffer (Promega) and Luciferase activity was measured as described above.

### RT-PCR and Quantitative PCR

Total RNA was extracted from different cell types or tissue using RNAeasy Plus extraction kit (Qiagen). For RNA extraction from human kidney, tissues were frozen in liquid nitrogen and stored at −80C. 30 mg of tissue was homogenized in a bullet blender (Next Advance) in buffer RLT Plus (Qiagen) and RNA was extracted as described above. For RT-PCR, 2 µg of total RNA was reverse transcribed into cDNA using random primers and MML-V reverse transcriptase (Promega) according to the manufacturer’s instruction. The PCR was performed with Pfu Ultra for cloning or Taq polymerase for analytical purposes respectively. MMP-1 primers have been described (Guo et al., 2006). The following primer set was used for the identification of Stx3S, forward hStx3Sxon9_for 5′-GGAGAAGGCACGAGATGAAA-3′ and reverse hStx3Sxon11_rev 5′-GGTTGCAAGGAAACAAAGGA-3′. Stx3S was cloned from HEK293T cells using two different primer sets. Set1 was used for the initial amplification from cDNA as a template using the forward primer hStx3C_F 5′-ATGAAGGACCGTCTGGAGCAGCTG-3′ and the reverse primer hStx3C_R 5′-TCATCTGAAGACAAGGGTGGCCAC-3′. The re-amplification was performed with the following second set of primers hStx3C_EcoRI_for_1 5′-TTTAAAAAGAATTCATGAAGGACCGTCTGGAGCAGCTG-3′ and hStx3_NotI_rev_1 5′-TTTTTAAAGCGGCCGCTCTGAAGACAAGGGTGGCCACACC-3′ and the PCR product was ligated into pcDNA4/TO/2xmyc/His. Stratagene Mx3000P Real-Time system (Agilent Technologies, Santa Clara, CA) were used for the qPCR reaction. In addition to the aforementioned MMP-1 primer set, the following primers were used for qPCR: FNTA, fwd 5′ - GCAGGATCGTGGTCTTTCCA - 3′, rev 5′ - AACTCCTGCAGGGTCCCTTA - 3′; KDM2B fwd 5′-GACTTGTCGGACGTGGAGGA - 3′, rev 5′ - CACATGTTGTCCACCCAGTC - 3′ (Vlahopoulos et al., 2008); SART3, fwd 5′ - TCAATGGCAAGAAAACGGGT - 3′, rev 5′ - AAGCCATGGCGTACTCATCC - 3′; GAPDH, fwd 5′ - AGCAAGAGCACAAGAGGAAGAG - 3′, rev 5′ - GAGCACAGGGTACTTTATTGATGG - 3′; β-actin, fwd 5′ - GAAGTGTGACGTTGACATCC - 3′, rev 5′ - ACAGAGTACTTGCGCTCAGG - 3′. Relative quantitation using the 2^(-∆∆Ct)^ method was used to determine gene expression.

### Microarray gene expression analysis

Microarray analysis of Caco2 cells stably expressing shRNA against Stx3S was set up in triplicate. Briefly, cells were lysed and total RNA was isolated using a RNeasy kit from Qiagen (Valencia, CA). 5µg of RNA were used to produce cDNA followed by generation of biotinylated cRNA. Samples were analyzed by the University of Pennsylvania School of Medicine Microarray facility by hybridization to GeneChip Human Gene 2.0 ST arrays (Affymetrix, Santa Clara, CA). Comparisons between group means were done by a two-sided heteroscedastic Student′s t-test.

### Viability/proliferation

In quadruplicate, Caco2 cells stably expressing shRNA against Stx3S were seeded in complete media. After 24h, media was changed to serum-free or complete media. At indicated time points after changing media, wells were analyzed using the CellTiter-Glo Luminescent Cell Viability Assay (Promega, Madison WI) per manufacturer’s instructions. Statistical analysis was performed in Prism4 (GraphPad Software, La Jolla, CA).

In triplicate, Caco2 cells stably expressing shRNA against Stx3S or scrambled control were seeded in complete media. Media was changed 24h after seeding. Five days after media change, cells were harvested and processed as above for immunocytochemistry. Cyclin B1 and cleaved Caspase-3 positive cells were counted using FIJI software. An average of 250 cells were counted per replicate. Statistical analysis was performed in Prism4 (GraphPad Software, La Jolla, CA).

## ACKNOWLEDGMENTS

We thank Roger Davis (University of Massachusetts), John Hassell (McMaster University) and Andrew Sharrocks (University of Manchester) for reagents and advice. This work was supported by grants from the NIH (DK62338), and the California Cancer Research Coordinating Committee to T.W., and Postdoctoral Fellowships from the Cancer Center of Santa Barbara and the Spanish Ministry of Education and Science to C.W. and E.R., respectively. J.B.H. is supported by 1R01NS054794 from NINDS, and 1R01HL097800 from NHBLI.

## REFERENCES

Amelio, A.L., L.J. Miraglia, J.J. Conkright, B.A. Mercer, S. Batalov, V. Cavett, A.P. Orth, J. Busby, J.B. Hogenesch, and M.D. Conkright. 2007. A coactivator trap identifies NONO (p54nrb) as a component of the cAMP-signaling pathway. Proc Natl Acad Sci U S A. 104:20314–20319.

Bock, J.B., H.T. Matern, A.A. Peden, and R.H. Scheller. 2001. A genomic perspective on membrane compartment organization. Nature. 409:839–841.

Bosc, D.G., B.S. Goueli, and R. Janknecht. 2001. HER2/Neu-mediated activation of the ETS transcription factor ER81 and its target gene MMP-1. Oncogene. 20:6215–6224.

de Launoit, Y., J.L. Baert, A. Chotteau-Lelievre, D. Monte, L. Coutte, S. Mauen, V. Firlej, C. Degerny, and K. Verreman. 2006. The Ets transcription factors of the PEA3 group: transcriptional regulators in metastasis. Biochimica et biophysica acta. 1766:79–87.

Deane, R., W. Schafer, H.P. Zimmermann, L. Mueller, D. Gorlich, S. Prehn, H. Ponstingl, and F.R. Bischoff. 1997. Ran-binding protein 5 (RanBP5) is related to the nuclear transport factor importin-beta but interacts differently with RanBP1. Molecular and cellular biology. 17:5087–5096.

Delgrossi, M.H., L. Breuza, C. Mirre, P. Chavrier, and A. Le Bivic. 1997. Human syntaxin 3 is localized apically in human intestinal cells. Journal of cell science. 110:2207–2214.

Dulubova, I., S. Sugita, S. Hill, M. Hosaka, I. Fernandez, T.C. Südhof, and J. Rizo. 1999. A conformational switch in syntaxin during exocytosis: role of munc18. The EMBO Journal. 18:4372–4382.

Eswaran, J., A. Horvath, S. Godbole, S.D. Reddy, P. Mudvari, K. Ohshiro, D. Cyanam, S. Nair, S.A. Fuqua, K. Polyak, L.D. Florea, and R. Kumar. 2013. RNA sequencing of cancer reveals novel splicing alterations. Scientific reports. 3:1689.

Fasshauer, D., R.B. Sutton, A.T. Brunger, and R. Jahn. 1998. Conserved structural features of the synaptic fusion complex: SNARE proteins reclassified as Q-and R-SNAREs. Proc Natl Acad Sci U S A. 95:15781–15786.

Gozdecka, M., and W. Breitwieser. 2012. The roles of ATF2 (activating transcription factor 2) in tumorigenesis. Biochemical Society transactions. 40:230–234.

Guo, B., R.E. Sallis, A. Greenall, M.M. Petit, E. Jansen, L. Young, W.J. Van de Ven, and A.D. Sharrocks. 2006. The LIM domain protein LPP is a coactivator for the ETS domain transcription factor PEA3. Mol Cell Biol. 26:4529–4538.

Hong, W. 2005. SNAREs and traffic. Biochim Biophys Acta. 1744:120–144.

Horvath, A., S.B. Pakala, P. Mudvari, S.D. Reddy, K. Ohshiro, S. Casimiro, R. Pires, S.A. Fuqua, M. Toi, L. Costa, S.S. Nair, S. Sukumar, and R. Kumar. 2013. Novel insights into breast cancer genetic variance through RNA sequencing. Scientific reports. 3:2256.

Ibaraki, K., H.P. Horikawa, T. Morita, H. Mori, K. Sakimura, M. Mishina, H. Saisu, and T. Abe. 1995. Identification of four different forms of syntaxin 3. Biochemical and biophysical research communications. 211:997–1005.

Jagadish, M.N., J.T. Tellam, S.L. Macaulay, K.H. Gough, D.E. James, and C.W. Ward. 1997. Novel isoform of syntaxin 1 is expressed in mammalian cells. Biochem J. 321:151–156.

Keld, R., B. Guo, P. Downey, C. Gulmann, Y.S. Ang, and A.D. Sharrocks. 2010. The ERK MAP kinase-PEA3/ETV4-MMP-1 axis is operative in oesophageal adenocarcinoma. Molecular Cancer. 9:313–313.

Kent, W.J., C.W. Sugnet, T.S. Furey, K.M. Roskin, T.H. Pringle, A.M. Zahler, and D. Haussler. 2002. The human genome browser at UCSC. Genome research. 12:996–1006.

Kottakis, F., P. Foltopoulou, I. Sanidas, P. Keller, A. Wronski, B.T. Dake, S.A. Ezell, Z. Shen, S.P. Naber, P.W. Hinds, E. McNiel, C. Kuperwasser, and P.N. Tsichlis. 2014. NDY1/KDM2B functions as a master regulator of polycomb complexes and controls self-renewal of breast cancer stem cells. Cancer research. 74:3935–3946.

Kreitzer, G., J. Schmoranzer, S.H. Low, X. Li, Y. Gan, T. Weimbs, S.M. Simon, and E. Rodriguez-Boulan. 2003. Three-dimensional analysis of post-Golgi carrier exocytosis in epithelial cells. Nat Cell Biol. 5:126–136.

Kuure, S., X. Chi, B. Lu, and F. Costantini. 2010. The transcription factors Etv4 and Etv5 mediate formation of the ureteric bud tip domain during kidney development. Development. 137:1975–1979.

Li, X., S.H. Low, M. Miura, and T. Weimbs. 2002. SNARE expression and localization in renal epithelial cells suggest mechanism for variability of trafficking phenotypes. Am J Physiol Renal Physiol. 283:F1111–1122.

Low, S.H., S.J. Chapin, T. Weimbs, L.G. Komuves, M.K. Bennett, and K.E. Mostov. 1996. Differential localization of syntaxin isoforms in polarized Madin-Darby canine kidney cells. Molecular biology of the cell. 7:2007–2018.

Low, S.H., S.J. Chapin, C. Wimmer, S.W. Whiteheart, L.G. Komuves, K.E. Mostov, and T. Weimbs. 1998. The SNARE machinery is involved in apical plasma membrane trafficking in MDCK cells. J Cell Biol. 141:1503–1513.

Low, S.H., X. Li, M. Miura, N. Kudo, B. Quinones, and T. Weimbs. 2003a. Syntaxin 2 and endobrevin are required for the terminal step of cytokinesis in mammalian cells. Dev Cell. 4:753–759.

Low, S.H., X. Li, M. Miura, N. Kudo, B. Quiñones, and T. Weimbs. 2003b. Syntaxin 2 and endobrevin are required for the terminal step of cytokinesis in mammalian cells. Developmental cell. 4:753–759.

Low, S.H., L.Y. Marmorstein, M. Miura, X. Li, N. Kudo, A.D. Marmorstein, and T. Weimbs. 2002. Retinal pigment epithelial cells exhibit unique expression and localization of plasma membrane syntaxins which may contribute to their trafficking phenotype. Journal of cell science. 115:4545–4553.

Low, S.H., M. Miura, P.A. Roche, A.C. Valdez, K.E. Mostov, and T. Weimbs. 2000. Intracellular redirection of plasma membrane trafficking after loss of epithelial cell polarity. Molecular biology of the cell. 11:3045–3060.

Lu, B.C., C. Cebrian, X. Chi, S. Kuure, R. Kuo, C.M. Bates, S. Arber, J. Hassell, L. MacNeil, M. Hoshi, S. Jain, N. Asai, M. Takahashi, K.M. Schmidt-Ott, J. Barasch, V. D'Agati, and F. Costantini. 2009. Etv4 and Etv5 are required downstream of GDNF and Ret for kidney branching morphogenesis. Nature genetics. 41:1295–1302.

Martin-Martin, B., S.M. Nabokina, P.A. Lazo, and F. Mollinedo. 1999. Co-expression of several human syntaxin genes in neutrophils and differentiating HL-60 cells: variant isoforms and detection of syntaxin 1. J Leukoc Biol. 65:397–406.

Nakayama, T., H. Kamiguchi, and K. Akagawa. 2012. Syntaxin 1C, a soluble form of syntaxin, attenuates membrane recycling by destabilizing microtubules. Journal of cell science. 125:817–830.

Nakayama, T., K. Mikoshiba, T. Yamamori, and K. Akagawa. 2003. Expression of syntaxin 1C, an alternative splice variant of HPC-1/syntaxin 1A, is enhanced by phorbol-ester stimulation in astroglioma: participation of the PKC signaling pathway. FEBS Lett. 536:209–214.

Nakayama, T., K. Mikoshiba, T. Yamamori, and K. Akagawa. 2004. Activation of syntaxin 1C, an alternative splice variant of HPC-1/syntaxin 1A, by phorbol 12-myristate 13-acetate (PMA) suppresses glucose transport into astroglioma cells via the glucose transporter-1 (GLUT-1). The Journal of biological chemistry. 279:23728–23739.

Nishida, T., M. Terashima, K. Fukami, and Y. Yamada. 2007. Repression of E1AF transcriptional activity by sumoylation and PIASy. Biochem Biophys Res Commun. 360:226–232.

O'Hagan, R.C., R.G. Tozer, M. Symons, F. McCormick, and J.A. Hassell. 1996. The activity of the Ets transcription factor PEA3 is regulated by two distinct MAPK cascades. Oncogene. 13:1323–1333.

Pereira, S., A. Massacrier, P. Roll, A. Verine, M. Etiennegrimaldi, Y. Poitelon, A. Robagliaschlupp, S. Jamali, N. Roeckeltrevisiol, and B. Royer. 2008. Nuclear localization of a novel human syntaxin 1B isoform. Gene. 423:160–171.

Perlmann, T., and L. Jansson. 1995. A novel pathway for vitamin A signaling mediated by RXR heterodimerization with NGFI-B and NURR1. Genes Dev. 9:769–782.

Peterson, M.D., and M.S. Mooseker. 1992. Characterization of the enterocyte-like brush border cytoskeleton of the C2BBe clones of the human intestinal cell line, Caco-2. Journal of cell science. 102:581–600.

Quinones, B., K. Riento, V.M. Olkkonen, S. Hardy, and M.K. Bennett. 1999. Syntaxin 2 splice variants exhibit differential expression patterns, biochemical properties and subcellular localizations. Journal of cell science. 112:4291–4304.

Rodriguez-Boulan, E., and W.J. Nelson. 1989. Morphogenesis of the polarized epithelial cell phenotype. Science. 245:718–725.

Rothman, J.E. 2014. The principle of membrane fusion in the cell (Nobel lecture). Angew Chem Int Ed Engl. 53:12676–12694.

Sharma, N., S.H. Low, S. Misra, B. Pallavi, and T. Weimbs. 2006. Apical targeting of syntaxin 3 is essential for epithelial cell polarity. J Cell Biol. 173:937–948.

Singha, P.K., S. Pandeswara, M.A. Venkatachalam, and P. Saikumar. 2013. Manumycin A inhibits triple-negative breast cancer growth through LC3-mediated cytoplasmic vacuolation death. Cell death & disease. 4:e457.

Sudhof, T.C. 2014. The molecular machinery of neurotransmitter release (Nobel lecture). Angew Chem Int Ed Engl. 53:12696–12717.

Sutton, R.B., D. Fasshauer, R. Jahn, and A.T. Brunger. 1998. Crystal structure of a SNARE complex involved in synaptic exocytosis at 2.4 A resolution. Nature. 395:347–353.

Talbot, J.J., J.M. Shillingford, S. Vasanth, N. Doerr, S. Mukherjee, M.T. Kinter, T. Watnick, and T. Weimbs. 2011. Polycystin-1 regulates STAT activity by a dual mechanism. Proceedings of the National Academy of Sciences of the United States of America. 108:7985–7990.

Vecchi, M., S. Polo, V. Poupon, J.W. van de Loo, A. Benmerah, and P.P. Di Fiore. 2001. Nucleocytoplasmic shuttling of endocytic proteins. The Journal of cell biology. 153:1511–1517.

Vlahopoulos, S.A., S. Logotheti, D. Mikas, A. Giarika, V. Gorgoulis, and V. Zoumpourlis. 2008. The role of ATF-2 in oncogenesis. Bioessays. 30:314–327.

Weimbs, T., S.H. Low, S.J. Chapin, and K.E. Mostov. 1997a. Apical targeting in polarized epithelial cells: there's more afloat than rafts. Trends in cell biology. 7:393–399.

Weimbs, T., S.H. Low, S.J. Chapin, K.E. Mostov, P. Bucher, and K. Hofmann. 1997b. A conserved domain is present in different families of vesicular fusion proteins: a new superfamily. Proc Natl Acad Sci U S A. 94:3046–3051.

Weimbs, T., S.H. Low, X. Li, and G. Kreitzer. 2003. SNAREs and epithelial cells. Methods. 30:191–197.

Weimbs, T., K. Mostov, S.H. Low, and K. Hofmann. 1998. A model for structural similarity between different SNARE complexes based on sequence relationships. Trends in cell biology. 8:260–262.

Wickner, W., and R. Schekman. 2008. Membrane fusion. Nat Struct Mol Biol. 15:658–664.

Zhang, Q. 2005. The Subcellular Localization of Syntaxin 17 Varies Among Different Cell Types and Is Altered in Some Malignant Cells. Journal of Histochemistry and Cytochemistry. 53:1371–1382.

